# The tempest in a cubic millimeter: Image-based refinements necessitate the reconstruction of 3D microvasculature from a large series of damaged alternately-stained histological sections

**DOI:** 10.1101/2019.12.17.877738

**Authors:** Oleg Lobachev

## Abstract

This work presents two methods that facilitate a 3D reconstruction of microscopic blood vessels in the volume slightly larger than 1 mm^3^. The source of the data are histological serial sections, i.e., microscopic images of probes, stained with immunohistochemistry. Odd and even sections have different stainings in our primary data set. Thus, firstly, an approach to register an alternately-stained series is presented. With image filtering and a feature-detection-based registration we obtain a registered stack of 148 serial sections. The series has missing sections, locally damaged sections, artifacts from acquisition. All these hinder correct connectivity of blood vessels. With our second approach we interpolate the missing information while maintaining the connectivity. We achieve this with deformations based on dense optical flow. The presented methodology is applicable to further histological series. A combination of both approaches allows us to reconstruct more than 76% larger volumes. An important detail was the composition mode of images. Summarizing, we use methods from image processing and computer vision to create large-scale 3D models from immunostained histological serial sections.

## I. INTRODUCTION

Histological serial sections are the method of choice for obtaining insights from human tissues at microscopic levels. Despite the availability of further imaging techniques, ranging from micro-CT to two-photon microscopy, there is de-facto no choice when working with human specimens. Micro-CT lacks on resolution and does not allow for labeling of specific cells. Nano-CT has too small working area. Virtually all recently developed imaging methods focus on model organisms. In them, artificial enhancements are possible, including fluorescent proteins, artificially-transparent tissues (incl. “clarity”), and further modifications that facilitate novel imaging methods. Those cannot be thought of with humans; the specimens need to be processed as they are. Immunohistology introduced a much more precise staining. With immunohistology, specific molecules can be visualized by “connecting” them to the antibodies that in their turn make the color pigment solution precipitate where the sought-for molecules were in the first place. Those molecules are not necessarily unique, but in most cases, morphological differences between different cell types exhibiting the same molecule is large enough for a proper diagnosis. For an introduction to histology, consult, e.g., ref. [58].

A single histological section, even with immunostaining, does not suffice for a proper understanding. The sections are very thin (typically about 7 μm in standard sections, starting from 1 μm in semi-thin sections), but cover a larger area (about 1 cm^2^). This yields a de-facto two-dimensional representation of the tissue. In longer-spawning entities, such as blood vessels or nerves, very little insight can be derived from a single section. Fortunately, a series of histological sections, the so-called serial sections, can be sectioned, stained, and processed.

The best way of facilitating better understanding from serial sections is to perform a full-fledged 3D reconstruction [66]. The path to it is, however, long and problematic. A proper registration of the serial sections needs to be established [68]. The sections can vary in thickness and hence in intensity. They need to be normalized [53]. In further processing, a correct separation of colors in the staining [78], faithful selection of the shape, correct mesh construction [69] and mesh processing should follow. The presentation of the final reconstruction to the experts is also an issue. The interaction of the experts with the model, the so-called visual analytics, is the actual way of finding insights and producing results. We developed a virtual realty application for this sake [66, 99, 101].

This paper, however, focuses on the cases when not everything works well in the required biomedical processing and the above 3D reconstruction pipeline. The reasons vary greatly, but in many cases the result is the same: A section or a part thereof is not available for the 3D reconstruction.

Previously, the only definite way of coping with such problems was to truncate the series. While this is the most pure and not disputable approach, the amount of data to be discarded is quite extreme. In the series we mostly focus on this paper, 150 sections were produced, first and last were controls and were not indented for the 3D reconstruction. Due to problems during biological processing, 25 sections in the beginning of the series could not be fully used. Two sections were outright lost. Acquisition of such a large data set is a tedious and error-prone process. Because of some focusing problems, not all section images were usable. When the series was truncated to mitigate all above problems, only 83 to 91 sections were usable in varying regions. This constitutes up to 44.67% of lost sections. (Fig. 1 presents a truncated and a healed mesh.) With the methods presented here, we were able to salvage the damaged regions and to bridge over the lost sections, reconstructing all 148 immunostained sections. We operate on a region of interest (ROI) of 2*k* × 2*k* pixels, yielding a 3D reconstruction of more than 1 mm^3^ of human splenic tissue at acquisition resolution of 0.5 μm/pixel.

**Figure 1:**
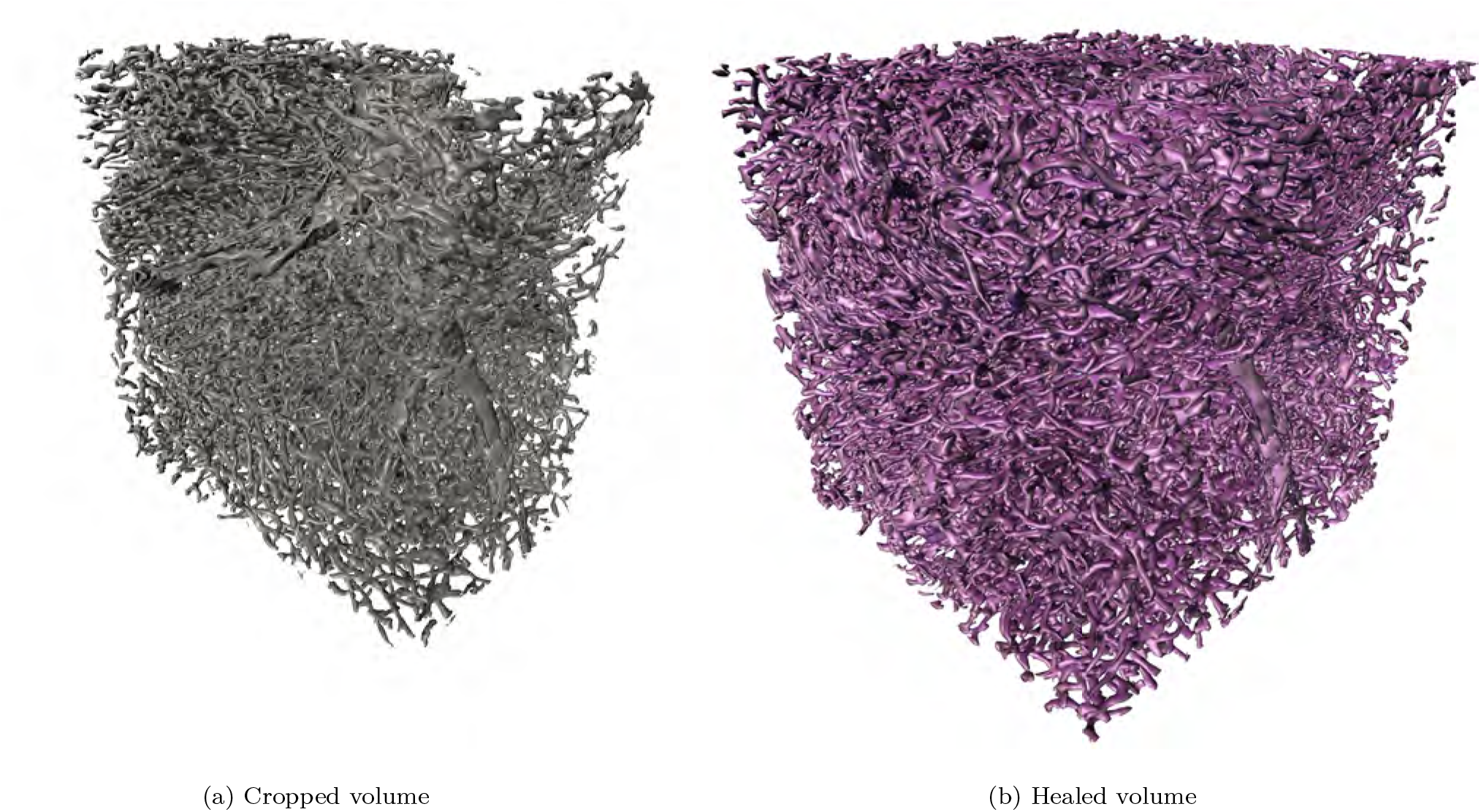
Motivation for this paper: healing greatly increases the amount of information available for analysis. Here, two decimated meshes are presented, which were reconstructed at same iso-value 120 from corresponding volumes. The meshes are rendered from the same view point. Fig. (a) shows a reconstruction from 84 sections available in the current region without healing. In (b) the result of our healing method is presented, all 148 immunostained sections are used. Notice also the high degree of complexity and self-occlusion. Fig. 9 compares the full meshes. The *x* and *y* sides of the reconstructions (the wider sides in (a)) span over 1 mm.

### A. Related Work

There is a lot of related work on registration, including co-registration of different stainings (Section IA1) and co-registration of other modalities (Section I A 2). There is virtually no related work on the repair of histological sections (Section I A 3). We also present an overview of related deep learning approaches (Section I A 4) and mention the methods we use for evaluation in Section I A 5.

#### 1. Registration of serial sections

Ourselin et al. [80] presented one of the first fully automatic approaches for the “normal” registration of serial sections. Pichat et al. [84] feature a recent review.

In this work we utilize our feature-based non-rigid method [68, 106] for the actual registration of the whole series, extending it to work with alternating staining. In has been also previously used for a fine-grain registration. For a pairwise deformation of neighboring sections, we also used `elastix` [55].

Kajihara et al. present a framework for registration of serial sections [51]. They perform a RANSAC-based rigid registration based on feature detection first, and then an as-rigid-as-possible region-based registration from keypoint correspondences is used. The regions emerge from clustering keypoints. Then, local transforms are blended together to obtain a non-rigid transformation. (In a contrast, our registration method [68] uses keypoint detection and matching; then RANSAC for the rigid phase; then non-rigid alignment is hierarchically computed from the neighborhood matches for the whole series of sections, producing a global pyramid of non-rigid transformations.) Since Kajihara et al. are concerned with larger series, a sudden discontinuity in registration is mitigated by manual selection of a reference. Our work is focused on an automatic registration (with possible manual removal of an offending image yielding a discontinuity) and, mostly important, the repair of the damage caused by such a removal or by inevitable processing and acquisition artifacts.

The remaining part of this section handles coregistration, i.e., the registration of differently stained sections. Refs. [19, 56] rely on the segmentation of “common ground” between the different stainings. Similarly do Wentzensen et al. [110] and Braumann et al. [18], as Wentzensen et al. register three different stainings to each other by the means of a segmentation. They applied some sophisticated sampling strategies and used a fuzzy classification of the segmented stainings. We recovered our similarity information in a pretty different manner.

With a similar goals Mueller et al. [77] focus on deformable registration of whole routine sections for digital pathology, involving `elastix` [55] and mutual information with additional constraints. Our work focuses on immuno-histological stainings.

Mosaliganti et al. [76] developed a registration method for larger stacks and were bound to introduce “a mixture of automated and semi-automated enhancements”. Our registration method is fully automated. User input was only used to determine the masks during“partial image” healing sessions.

Lippolis et al. [63] perform co-registration of biopsies or removed organs, combining H&E, immunohistochemistry, and a fluorescence marker. They use RANSAC-based rigid registration, we co-register two immunohistological stainings and perform also multiple non-rigid steps. du Bois d’Aische et al. [31] register micro to macro images in a complex framework. We focus on one magnification level.

Jiang et al. [48] hierarchically co-registered H&E and immunohistochemistry (Caspase3, KI67 and PHH3) with image-patch-based method, involving FFT followed by kernel density estimation. They obtained the ground truth for the evaluation from manual registration. We used a fully different method stack for the registration, our evaluation is based on similarity metrics and 3D reconstruction; however our images feature a different grade of similarity.

Song et al. [96] use unsupervised classification for a co-registration of differently-stained serial sections. We pursue a fully different approach as our alternating series have some “common ground”. Ref. [105] develops a tool for a more comfortable diagnosis with co-registered differently-stained sections. They use chamfer matching for the registration. This work focuses on alternating series and repair of missing sections. Our series is coregistered to form a common volume, our evaluation [100] focuses on the analysis of the 3D models in virtual reality. Section IV 1 a discusses the methodological difference between the related work and our method.

#### 2. Co-registration in other modalities

Ashburner and Friston [6] co-register neuroimages using affine transformation and clustering on segmented images. Another early attempt was performed by Andersson et al. [5].

Bardinet et al. [12] have co-registered some serial sections and MRI of human brain. The utilized sections were, however, visually quite similar to MRI cross-sections (and additionally processed to increase the similarity). A registration of serial sections to the block face images (photographs of the probe before this particular section was made) has been used [82, 92] as a proxy for the registration with coarser, but inherently 3D data, such as MRI.

Michálek et al. [74] co-registered matrix-assisted laser desorption/ionization mass spectrometry and confocal fluorescence microscopic images. Such images have drastically different resolutions. Lorenzen et al. [70] coregistered different MR modalities (i. e., data from inherently 3D acquisitions) of the same brain. Ourselin et al. [81] parallelized the co-registration of MRI and CT data with the goal of a clinical use.

Lee et al. [59] are concerned with learning a similarity measure from aligned 3D images and then apply it to a rigid multimodal registration. Tang et al. [102] have a similar idea, but perform a deformable registration on MRI and CT. Heinrich et al. [42] derive a modality-independent descriptor and use it in multimodal MRI and CT registration.

Wachinger and Navab [107] derive a similarly looking representation from various modalities of 3D images and use it for registration. Haber and Modersitzki [41] introduce a different image similarity metric for multimodal registration of inherently 3D medical images. Gehrung et al. [36] co-register optoacoustic tomography data with MRI. Becker et al. [13] co-register micro-CT with histological sections with chamfer matching. Such co-registrations have clinical impact, e.g., [8, 60, 90]. A separate topic is a groupwise image registration, typically performed on MRI data, [11, 14, 15, 44, 45, 86, 111].

The actual idea of a co-registration is much broader, it has been used, e. g., in synthetic aperture radars [61] or mass spectrometry [83].

#### 3. Repair of damaged series

The quite wide-spread problem of damaged sections is not often mentioned in the literature, as it is widely regarded to be a technical issue.

Song et al. [97] are aware of the missing sections problem, no repair is performed, but attempts to recover the missing information in order to maintain the registration; naturally, their method loses precision in such cases. Burton et al. [21] mention the missing sections problem; missing data is approximated by averaging nearest neighbor sections. In this work we perform a more elaborate repair, involving optical flow and a meaningful blending.

In case of a missing section, Saleem and Logothetis [91] sketch drawings from previously obtained MRI. In a contrast, we derive information from neighboring sections.

Alic et al. [2] are aware of possibly damaged sections, but do not elaborate. They co-register MR and histological images with larger (60 μm) distance between sections, using a manual and derived-from-manual segmentations. Their registration of serial sections uses mutual information. We use automatic key-point-based matching.

Generation of intermediate sections for 3D reconstruction from microscopic data has been suggested before [22, 67], however the method presented here differs in goal, details of blending, and in using registration prior to the computation of optical flow. A comparison with existing methods is in Section IV 1 b.

#### 4. Deep learning

Lai [57], Litjens et al. [64, 65], Zhou et al. [116], Ker et al. [52] and Altaf et al. [4] provide an overview of modern machine learning techniques in application to medical imaging. Garcia-Garcia et al. [35] provide a review of machine learning techniques w.r.t. general segmentation.

Further papers [23, 30, 34, 71, 75, 87, 104, 108, 112, 113] highlight specific applications of machine learning to medical segmentation or further (medical) image processing. Ref. [24] is a current result in semantic segmentation. Quite a few [25, 72, 73, 94] have utilized deep learning to perform a multimodal registration. Deep learning is also used for the separation of staining colors [3, 3, 20, 39, 93] or for registration [10, 27, 28].

We do not use any neural networks in this work, but utilize conservative image processing and computer vision repertoire.

#### 5. Evaluation

We use dense optical flow [32] (see [9] for a review), Jaccard measure [46], and SSIM [109] to visually quantify the quality of our healing (Section III 1). The actual assessment can only happen through the evaluation of the quality of resulting 3D reconstructions (Section III 3). We use geodesic distance computations on the mesh [26, 29] for such an evaluation. (A recent development in this area is, e.g., ref. [79].)

### B. Contributions

Firstly, we present a method to “lift” a registration method to alternating stainings of histological serial sections, in case there is a common ground between stainings. Secondly, we introduce a novel method to repair damaged or missing sections in the series. Our method is able to maintain the microvessel connectivity. Thus, the size of section series available for a 3D reconstruction is largely enhanced. In our running example, the amount of sections increased from 84 to 148, i.e., an increase of more than 76%. With the presented method we were able to reconstruct the vasculature in slightly over 1 mm^3^ of tissue at a microscopic resolution.

## II. METHODS

### A. Data

#### 1. Data origin and description of the data set

We were primarily working with 148 serial sections from the spleen of a 22-year-old male, acquired in 2000. The ethics of work with human materials fulfilled local regulations of the Philipps-University Marburg at the time of acquisition. The goal of the study was to investigate the capillary sheaths and the distribution of B-lymphocytes. The sections were *alternatively* stained for CD34 (cells in the walls of blood vessels, blue), SMA (some supporting structures and environments of arterioles, larger blood vessels, brown), and either CD271 (capillary sheaths and weakly some cells in the follicles, red) or CD20 (B-lymphocytes, red). Fig. 2 provides an overview.

**Figure 2:**
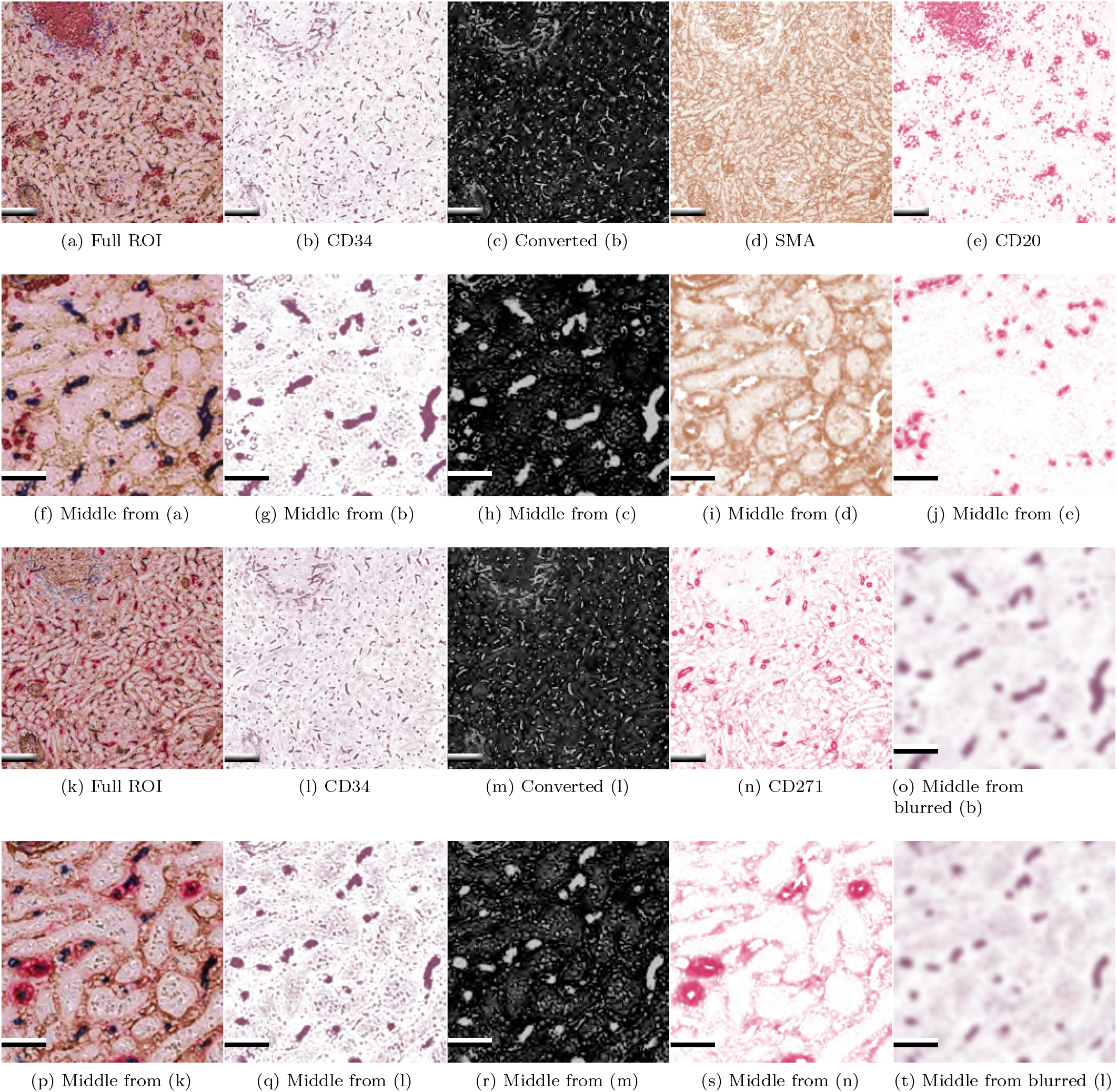
Overview of our main data set. The first and the third row show a 2.5*k* × 2.5*k* pixels ROI, the second and fourths row zoom in at the center to show details. Subfigures (a)–(j), (o) show registered section no. 10 (even, with red for B-lymphocytes), subfigures (k)–(n), (p)–(t) show registered section no. 11 (odd, with red for capillary sheaths). In all cases, blue pigment is present in CD34-positive cells, brown in SMA-positive. Subfigures (o), (t) show zoomed blurred CD34 channel processed in the same way, the registration input is. Subfigures (c), (h), (m), (r) show the input channel for further processing. Scale bars for full images (first row and (k)–(n)) are 200 μm, scale bars for center crops (second, fourth rows and (o)) are 50 μm.

The sections were processed for the transmitted light microscopy. This decision was made because of the established staining procedures and because the transmitted light stainings are permanent and can be stored and reexamined over decades.

The sections were digitized using Leica SCN 400 scanning microscope with a 20× lens. The final resolution was 0.5 μm/ pixel, full sections spanning approx. 8 mm × 11 mm were captured. A single section was 7 μm thick, hence the thickness of a full series of 148 immunostained sections covers the distance of more than 1 mm.

#### 2. Classification of damaged sections

Before we can continue with the description of the registration and of the repair measures, we need to distinguish between different kinds of damage in the digitized serial sections. Fig. 3a shows a non-damaged section, stained for CD34, SMA, CD271. Possible damage includes:

1. A physically missing section. The section in total was lost during biological processing.
2. Damaged part of a section. Some region in the section sustained damage, that is visible in the image regardless the acquisition kind, see Fig. 3b.
3. A larger part of a section or whole section is not in focus. Short of repeated acquisition, nothing can be done for this section, see Fig. 3c.
4. Some region of a section is not in focus or damaged in a different manner; however, parts of a section are in focus and can be successfully used for the section-wide registration, compare Figs. 3d, 3e.

**Figure 3:**
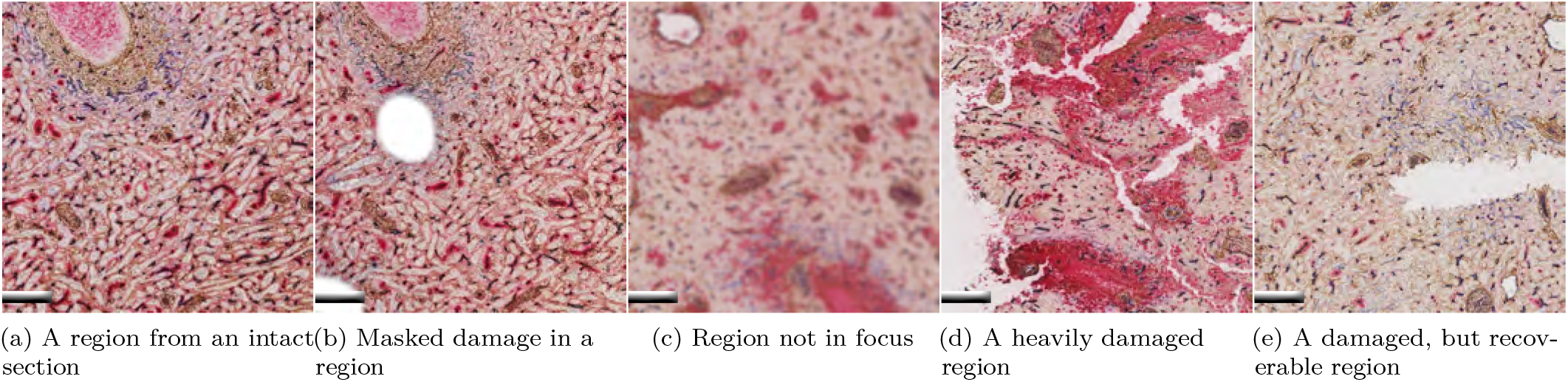
Examples for different kinds of damage in serial sections. All scale bars are 200 μm.

Boiled down to a ROI the above classification leads to two cases.

- An image in the series is irrecoverably lost and needs to be replaced.
- A part of an image in the series is lost, but other parts are usable.

### B. Processing

#### 1. Registration of alternating series

The methods for the registration of different stainings are based on either human input or on some degree of similarity between the different modalities. Machine learning from the annotations of identical or similar objects in varying stainings is able to identify “common” labels that are, in their turn, registered to each other. In our case, we were not able to afford the luxury of manual labeling. A mutual information or an extended machine learning approach on the “extracted” CD34 channels appears viable, but we chose a different approach.

The biological experiment was designed to have some common stainings between two subseries, the CD34 and SMA channels. A known method to separate the different colors in a histological staining is color deconvolution [78]. We aimed to apply our previously developed registration method [68] to alternating series. This method is based on feature detection, matching, and non-rigid deformations. The latter were computed with a Krylov subspace optimization method [33]. For the actual deformations, B-splines were used. We decoupled the “detection” images (the images the registration works on) from the “transformation” images (the images the established transformations are applied to). We found it crucial to denoise the feature detection inputs using Gaussian blur with *σ* = 6 (see also Figs. 2o, 2t). Without blurring, we found it impossible to register the series. We speculate that the noise visible in Figs. 2g, 2q produced a lot of small, very similar features. These confused the non-rigid registration.

We obtained a registration of a full series in its original resolution (thousands of pixels per side). For it, we used a blur-based denoising; the registration was applied to the full sections (mitigating local problems); instead of the missing sections and of some damaged sections duplicates of their neighbors were used as a placeholder. The latter issue necessitates further processing.

A rigid phase and 4 levels on non-rigid deformations, were established on the blurred images and subsequently applied to the actual full series of complete sections. In our experiments we used the CD34 (blue) channel for the registration, as this channel contains continuous structures (blood vessels). Acquisition artifacts in the series also influenced the registration. We stress that above section replacements can be tolerated as an intermediate step only, as a missing section unequivocally means interruptions of the capillaries.

At this stage, a region of interest can be selected. In a typical 3D reconstruction pipeline we select ROIs of size 2.5*k* × 2.5*k* pixels for the final output of 2*k* × 2*k* pixels after the final interpolation step.

#### 2. Healing of separate channels

We used standard method by Reinhard et al. [89] as implemented by Khan et al. [53]. for the normalization of the ROIs.

The healing is commenced on normalized channel-separated images. First, a color deconvolution [78] is performed on the final registered images. We obtain four channels: CD34, SMA, CD271, and CD20. Color space conversion is performed from color-paletted 8-bit images (originating from color deconvolution) to intensity-based single-channel 8-bit images. To give an example, in Fig. 2 we converted the output of color deconvolution for CD34 (b), (l) to a negated green RGB channel (c), (m).

##### a. Healing in general

We routinely use our optical-flow-based interpolation [67] to reduce the anisotropy of the data in our reconstructions [99, 101]. Having initially acquired the data with spatial resolution of 0.5 μm/voxel × 0.5 μm/voxel × 7 μm/voxel, we produce volumes with 0.5 μm/voxel × 0.5 μm/voxel × 1 μm/voxel for the actual 3D reconstruction. We heavily modify this interpolation mechanism for the healing of missing sections.

Due to the nature of CD20 labeling (B-cells), we do not need to perform a “connecting” interpolation on it. SMA is rather used as a guide to distinguish larger arterioles from smaller capillaries; it can sustain some damage or replicated sections. CD271 appears only in every other section, so interpolation is needed here. We also apply the below method to CD271, but it is not as crucial there as it is for CD34. Repeated sections or missing image parts produce interruptions in blood vessels that are reconstructed from CD34 labeling. Such interruptions change the topology and connectivity of the blood vessel network. Hence, CD34 channel has to be healed.

##### b. Healing a completely missing image

There is no “extra” information to come by, when an image is completely missing. Let us denote the image sequence in question with *A–X–B*. Image *X* is missing; images *A* and *B* are its neighbors. In some cases, to reduce the movement in the next step, we pairwise register *A* to *B*(yielding *A*′) and *B* to *A* (yielding *B*′) with `elastix` [55]. Notice that `elastix` is applied to already registered series. In other cases the movement can be bridged with optical flow, hence there we just set *A*′: = *A* and *B*′: = *B*.

Next, we create an optical-flow-based intermediate images from *A*′ to *B*′, resulting in the end with an image we call here *X*′. This way we aim to approximate the moving from *A* to *X* and from *X* to *B* in the whole series. The “movement” (originating from the registration and from the optical flow intermediates) is also of benefit for the 3D reconstruction, as it alleviates the artifacts from just mechanical replication of *A* + *B* as purported *X* (cf. Table I and Fig. 8e). With it, some capillary connections are maintained that would be broken otherwise (see also Fig. 9). Section IIB2d details on optical flow and intermediates. The nature of the composition operator + is a separate issue, see Section IIB 2 e.

**Table I:** Numerical measures for the images from Fig. 6. The final result of our method is *X*′, i.e., Fig. 6b The image *Z* is the result of our method on *A* and *B*, i.e., Fig. 6a. Basically, in the case of *Z* the registration is omitted. The images *T* and *T*′ are the result of maximum intensity blending applied to the inputs *A* and *B* and to registered inputs *A*′ and *B*′ correspondingly. They basically omit the optical flow from our method. The Jaccard measure was computed on images with binary threshold 120. Values range between 0 and 1. The higher the values the better.

| Image pair | SSIM | SSIM Figure | Jaccard |
| --- | --- | --- | --- |
| $A-B$ | 0.133 | 5c, 5g | 0.123 |
| $A'-B'$ | 0.0935 | 5k, 5o | 0.0100 |
| $A-A'$ | 0.159 | – | 0.0219 |
| $Z-X'$ | 0.143 | 6c, 6g | 0.359 |
| $A-X'$ | 0.105 | 6i, 6m | 0.238 |
| $X'-B$ | 0.112 | 6j, 6n | 0.251 |
| $T-X'$ | 0.118 | – | 0.342 |
| $T'-X'$ | 0.157 | – | 0.432 |
| consecutive | 0.185 | – | 0.285 |

##### c. Healing missing parts of an image

For the missing parts on an image the procedure is similar. Consider the same images *A, X*, *B* as above. However, as we now have a mask *M* on the damaged parts of the image *X*, instead of trying to “imagine” the whole contents of *X*, we need to do so only where *M* is not zero. The remaining, “valid” parts of *X* are mixed into the image *X*′. Also here the composition operator is important.

In the case of an out-of-focus image which is still marginally usable, the mask *M* is zero everywhere. The offending image is fittingly registered, because other regions outside of ROI are still in focus. In this case, adding it to the “imagined” image *X*′ enhances the result, as Fig. 4 shows.

**Figure 4:**
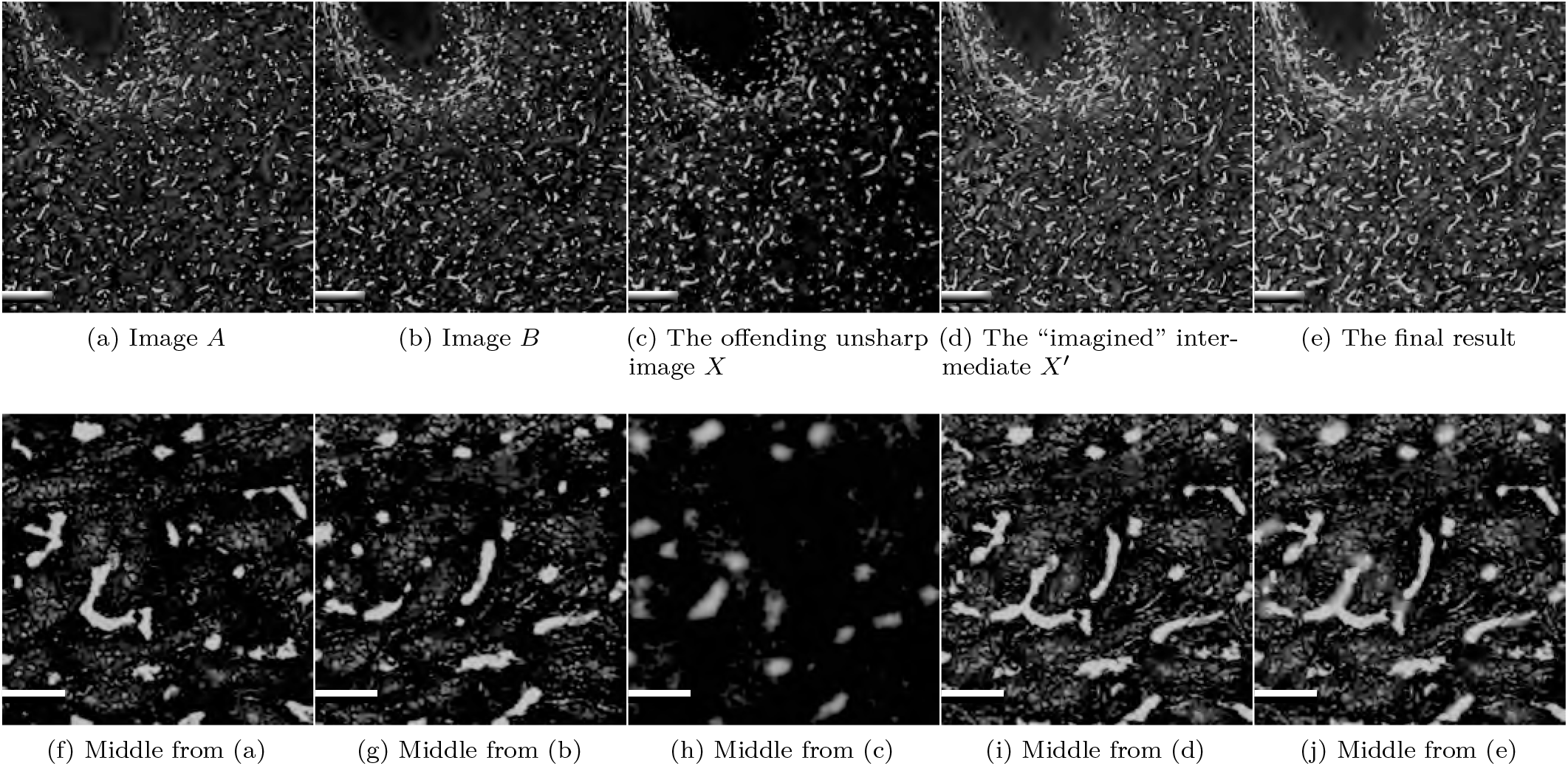
Healing an out-of-focus image. Still present information is reused. Scale bars in the top row are 200 μm, scale bars in the bottom row are 50 μm.

##### d. Optical flow and distortion

Dense optical flow by Farnebäck [32] models signals *f*_1_ and *f*_2_ as quadrics, e.g., *f*_1_(*x*) = *x^T^ A*_1_*x* + *b*_1_*x* + *c*_1_, where *A*_1_ is a symmetric matrix, *b*_1_ is a vector, and *c*_1_ is a scalar. The signal *f*_2_ has a similar representation, it is displaced by *d*. Local polynomial approximations are used below. We simplify the notation by writing *A* for a local approximation *A*(*x*).

Let *A*: = (*A*_1_ + *A*_2_)/2 and Δ*b*:= − (*b*_2_ – *b*_1_)/2. The distance between *f*_1_ and *f*_2_ can be viewed as a displacement field *d*, depending on *x*, with *Ad* = *b*. Let

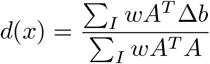

with sums running over all *x* in a region *I* and *w*(*x*) a weight for handling edges. In practice, the values *A^T^ A, A^T^* Δ*b*, Δ*b^T^* Δ*b* (for confidence) are computed point-wise. We use a multi-scale implementation from OpenCV [17,50].

Let 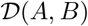 be the dense optical flow between matrices *A* and *B*. The flow is a vector field of local approximations *d*(*x*). We denote by 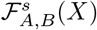 an operation than computes the flow 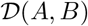 and then distorts *X* by 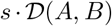 for a scale factor *s* ≥ 0. For example, 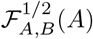 distorts *A* halfway to appear similar to *B*. We utilize OpenCV to apply the distortion.

##### e. Combining the flows for healing

In both above cases of healing we compute the optical flow 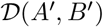 between *A*′ and *B*′ and also 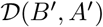. Then 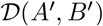 is used to distort *A*′ at the magnitude *s*, with 0 ≤ *s* ≤ 1, i.e., we compute 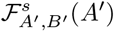. Similarly, we obtain the other direction 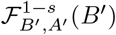. As we aim for a “middle ground”, we would combine 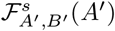 and 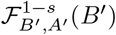 at *s* = 1/2. It is possible to use alpha-blending for the composition [67], but it is less useful here as Section III 3 reports. With alpha-blending, an intermediate is

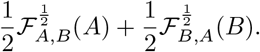

Crucial for the good results is the proper composition mode. Here we primarily use “lighten”, i.e.,

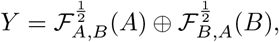

where ⊕ is the pixel-wise intensity maximum.

Finally, recall the presence of an image mask for healing only missing parts of an image. In this case, the nonmasked parts of *X* are combined with the image *Y* with ⊕, resulting in the final result *X*’.

### C. Further processing and visualization

After the healing, we interpolated the CD34, SMA, and CD271 volumes with our optical-flow-based method. The interpolated volumes were further processed in 3D Slicer [54, 85]—a bit differently for each channel. In case of CD34 (blood vessels), the volume processing consists of the morphological closing filter and Gaussian blurring. In some cases, e.g., for CD271, we also dilated the volumes. Then we generated the meshes [1, 7, 69] and processed them further [26, 49]. Section III 3 details on the mesh processing for this paper.

In practice, the final meshes were visualized in our virtual-reality-based application [66]. With the ability to place annotations and perform mesh painting, our software is used by the domain experts for knowledge discovery [99–101].

## III. RESULTS

With the presented method we were able to extend a consecutive series of 84 undamaged sections (Fig. 1a) to 148 sections spanning over a 1 mm in the direction of *z* axis (Fig. 1b). This means an grave 76.2% increase in number of available images. Section III 3 details on how the both above images were reconstructed.

In general, medical experts rated our healed meshes higher than the alternatives and similar in quality to the truncated meshes. The rating was based on experts’ perception of vasculature connectivity in ours reconstructions, which were inspected and quality controlled w.r.t. original registered sections in virtual reality. (Fig. 9 mimics such an analysis with geodesic distances.) Healing does not change intact images in the serial section stack. To quantify the difference between different methods, we use image- and mesh-based approaches below.

First, we evaluate our method with deformed and non-deformed input images, as well as compare the results of interpolation to the both neighbors in Section III 1. Sections III 2 a and III 2 b compare our healing method to ground truth on spleen and lung data sets. Section III 3 evaluates the 3D reconstructions from various healing strategies. The goal of this work was to restore broken connectivity in 3D reconstructions, hence it needs to be evaluated on those. All spleen images were normalized [89] to a common ground before processing.

### 1. Image-based evaluation of healing in production

Fig. 5 quantifies the `elastix`-based transformations of the input images. We compare original inputs *A* (a), (e) and *B* (b), (f) with *A*′ (*A* registered to *B*) (i), (m) and *B*′ (*B* registered to *A*) (j), (n) using SSIM (c), (g), (k),(o) and optical flow (d), (h), (l), (p).

**Figure 5:**
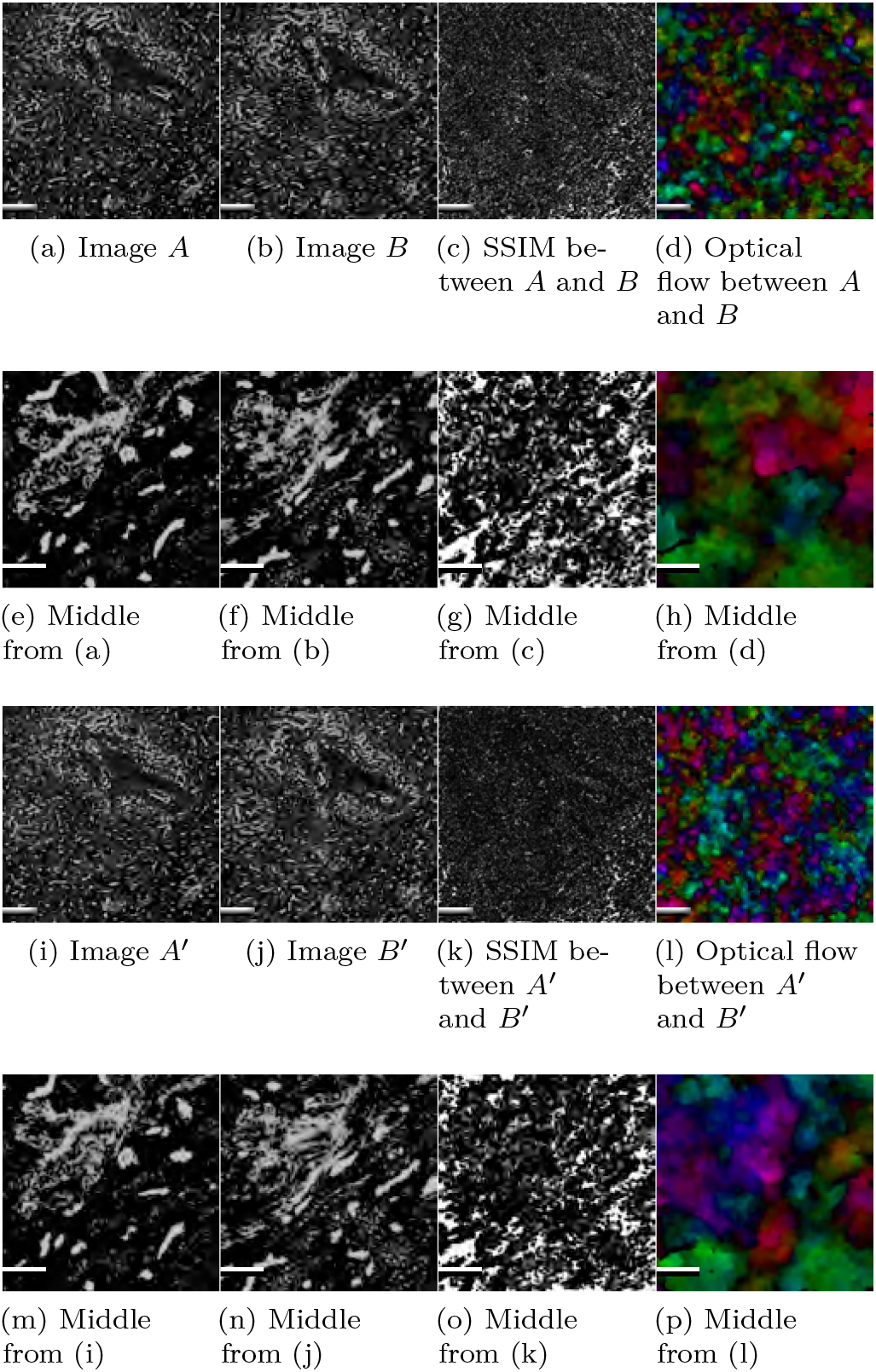
Normal and `elastix`-preregistered input for healing. Fig. 6 shows results. Table I presents SSIM and Jaccard values. Scale bars in the first and third rows are 200 μm, scale bars in the second and forth rows are 50 μm.

Fig. 6 showcases the results and quality measures of our method applied to the original inputs and to `elastix`-preregistered inputs. We see that SSIM is lower in *A*′–*X*′(i), (m) and *X*′–*B*′ (j), (n) pairs than in *Z–X*′ pair (c),(g), where *Z* is the result of our method without prior registration (a). Jaccard measure (Table I) supports this statement and optical flow visualization shows some local movement in Fig. 6 (d), (h), (k), (l), (o), (p). Optical flow visualizations appear similar between *A–B*, and *A–X*′, *X*′–*B* (Fig. 5, (l), (p), Fig. 6 (d), (h), (k), (l), (o), (p)), but consecutive sections yield less movement in optical flow (not shown). We argue that the local movement, shown to us by optical flow (k), (l), (o), (p), matches the desired outcome as we distort both images *A* and *B* to obtain a better intermediate.

**Figure 6:**
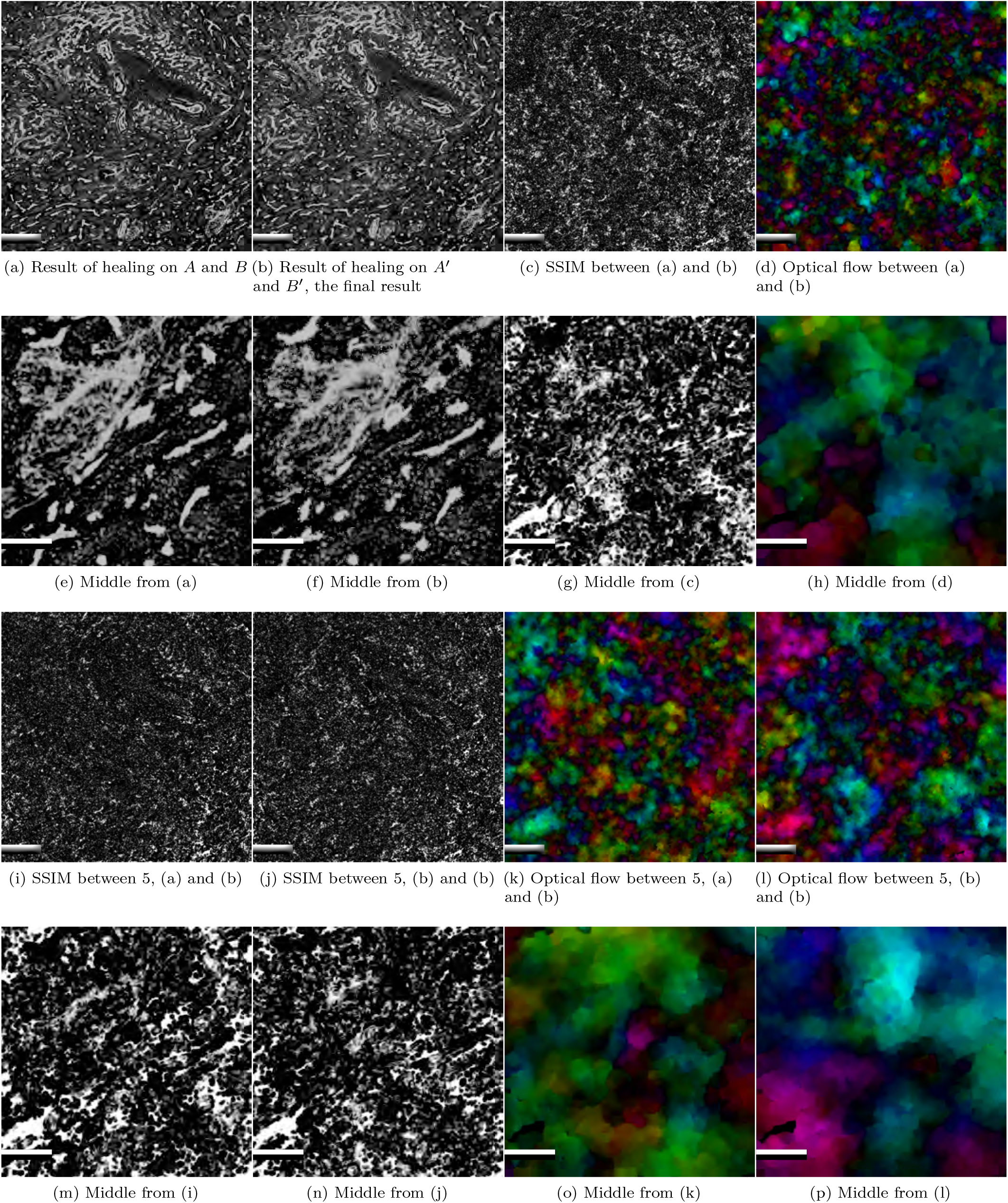
Healing in a case with `elastix`-preregistered input. Quantification of input images is in Fig. 5. See Table I for SSIM and Jaccard values. In SSIM, the brighter are the images, the better. In optical flow, the darker the images are, the better. Scale bars in the first and third rows are 200 μm, scale bars in the second and forth rows are 50 μm.

In Table I the differences in SSIM and Jaccard measure are similar between *A–X*′, *X*′–*B* and two consecutive sections. Surprisingly, both SSIM and Jaccard measure are very low for *A*′–*B*′ pair (Fig. 5k), i.e., for repeatedly registered input. A possible explanation is that *A*′ is distorted to fit *B*, but *B*′ is distorted to fit *A*. It is rather a way to distort both images towards an unavailable common ground *X* than a proper registration of *A* to *B*.

The highest Jaccard value results from an image pair *T*′–*X*′. Images *T* and *T*′ apply the blending method of our choice ⊕ to original (basically, *T* = *A* ⊕ *B*) and registered images (*T*′ = *A*′ ⊕ *B*′). Still, there is a substantial difference between *T*′ and *X*′. Similarly, there is a difference between *X*′ and *Z* (our method applied to *A* and *B*), resulting from repeated registration. Notice that our optical-flow-based interpolation also introduces local distortions.

### 2. Evaluation of healing on known images

#### a. Spleen

We took three undamaged consecutive sections from our main data set, cropped them to 2.3*k* × 2.3*k* pixels for more clarity, and pretended that the middle section needs to be healed. Full image was to be healed. We then subsequently computed the image measures similar to the previous section. No additional registration of images *A* and *B* was used. Fig. 7 and Table II report on our findings. The pair *A–X* is consecutive. SSIM between *X* and *X*′ (j), (n) appears lower (i.e., worse) than between *A* and *X*′ (i), (m). Optical flow visualizations are inconclusive in (l), (p) vs. (k), (o), but all those visualizations have some bright spots. Jaccard measure is slightly lower for *X–X*′. Summarizing, SSIM is higher at *A–X*′ pair, as *X*′ partially consists of *A*. In a contrast, Jaccard measure is similar between *A–X*′ and *X–X*′, the relatively high latter value allows us to rate the healing result as a success. Still, we evaluate also the models with different healing methods in Section III 3.

**Figure 7:**
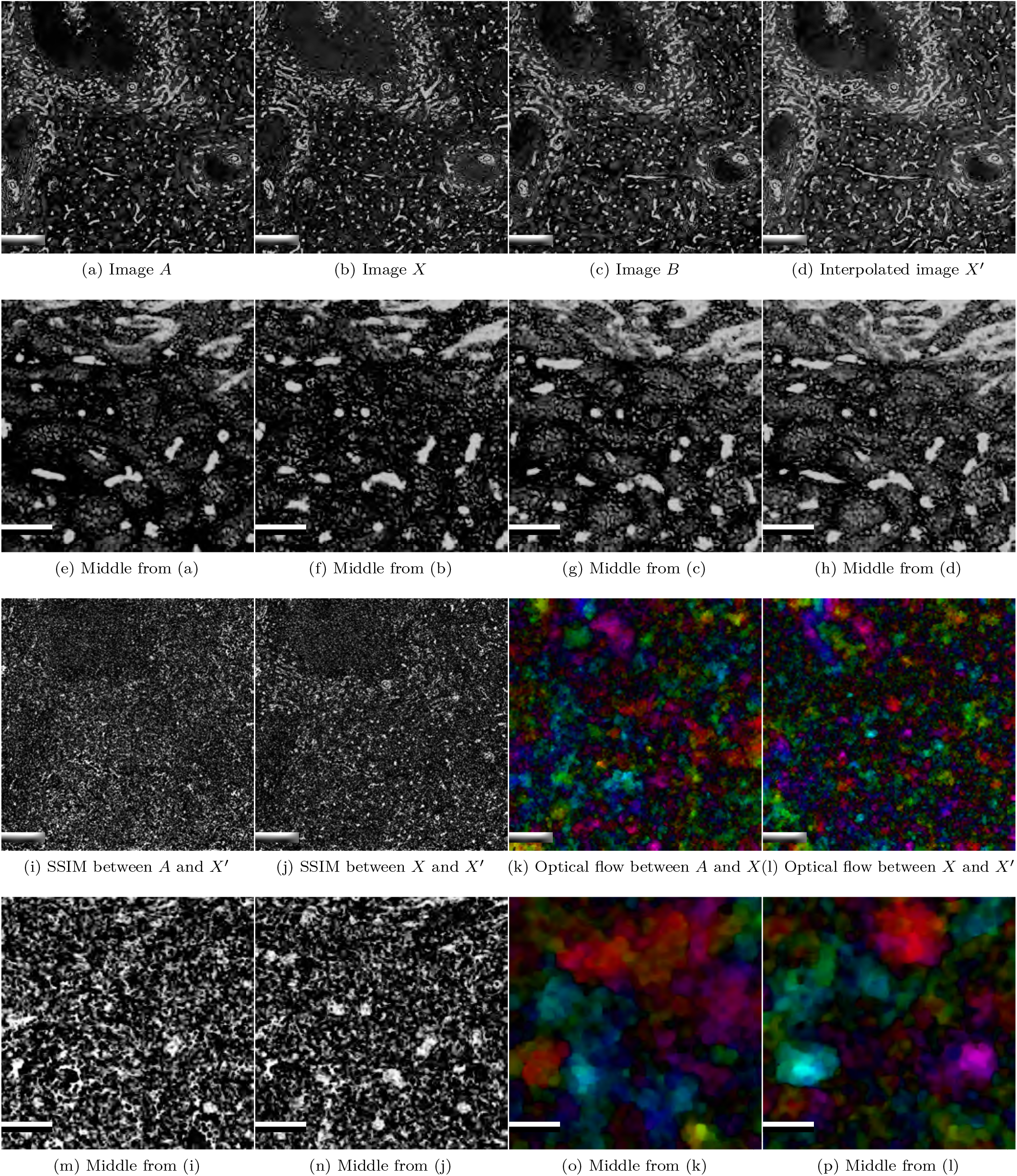
Evaluating our method with ground truth on a spleen data set. Image (b) is the ground truth, image (d) is the result of our method. In SSIM, the brighter are the images, the better. In optical flow, the darker the images are, the better. Scale bars in the first and third rows are 200 μm, scale bars in the second and forth rows are 50 μm.

**Table II:** Evaluating our method with ground truth on a spleen data set (Fig. 7). Image *A* precedes the actual interpolated image, image *X* is the ground truth, image *X*′ is the interpolated image. Values range between 0 and 1. The higher the values the better.

| Image pair | SSIM | Jaccard |
| --- | --- | --- |
| $A-X$ | 0.151 | 0.289 |
| $A-X'$ | 0.182 | 0.354 |
| $X-X'$ | 0.125 | 0.346 |

#### b. Lung

To demonstrate that our method works also on very different sections, we took a subseries of a rat lung serial sections [37, 38, 98]. Specifically, 2604 semithin sections with thickness of 1 μm of a pathological lung from a 12 week old Fischer F344 male rat were produced. The sections were stained with toluidine blue. A short resampled subseries with 800 × 800 pixels was selected for testing. It was rigidly pre-registered using our usual registration [68]. Images *A* and *B* were non-rigidly registered towards each other with `elastix` [55], resulting in *A*′ (a) and *B*′ (c). The images presented here were cropped after processing to 500 ×500 pixels to omit edge effects from registration. We did not apply normalization.

Fig. 8 shows the evaluation of our method on lung data with ground truth. SSIM and optical flow were computed on the full images, optical flow signaled larger changes at image edges. We show 500 × 500 center crops from those quality measures in Figs. (g), (h). SSIM between the ground truth *X* (b) and interpolated image *X*′ (d) on the full images, including the non-matching edges, is 0.846, SSIM between same cropped images is 0.935. We interpret this very high SSIM value as a success. Low values of the optical flow in Fig. 8h support this interpretation.

**Figure 8:**
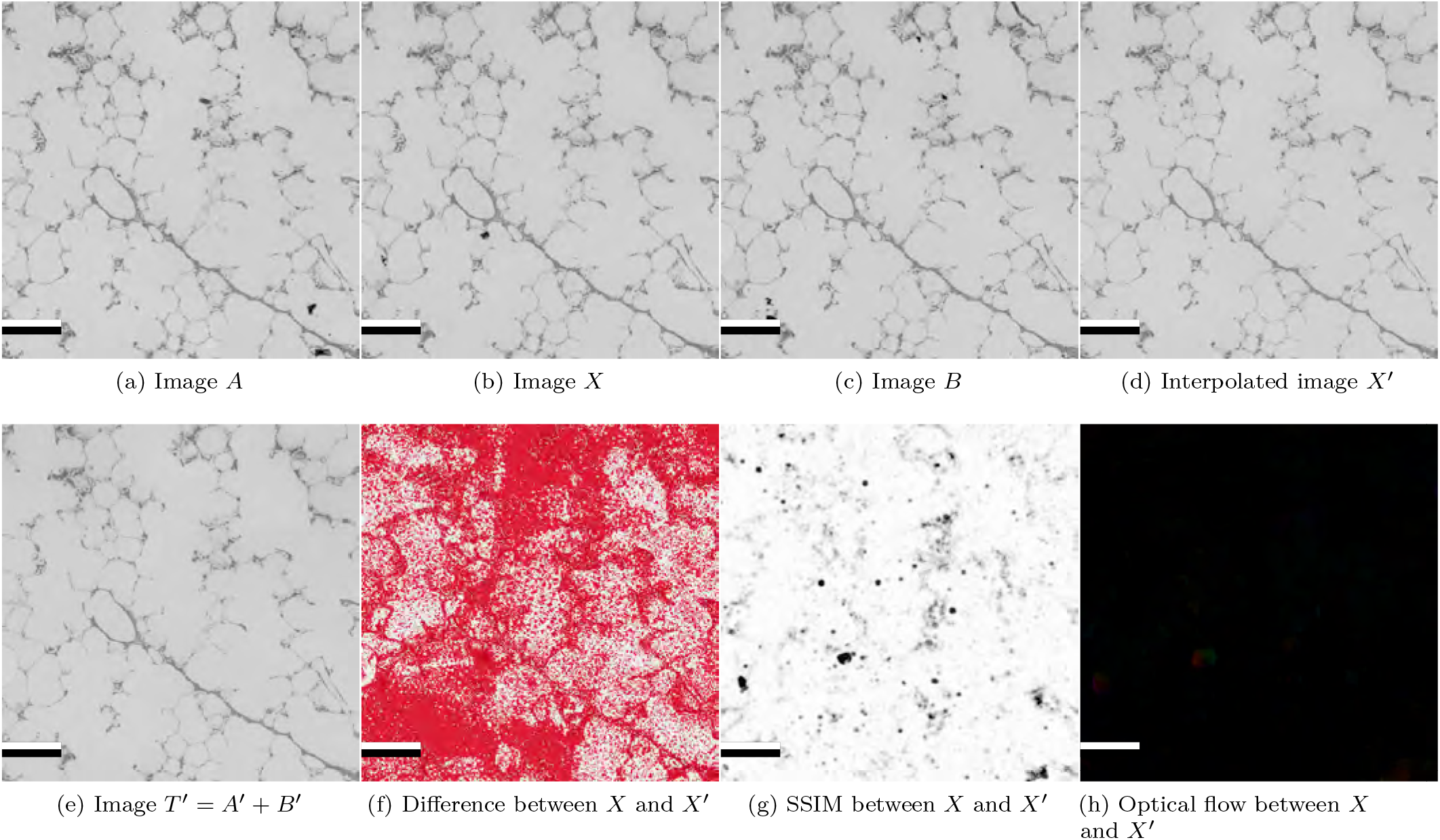
Evaluating our method with ground truth on a lung data set. Images *A* (a) and *B* (c) are used as an input. Image *X* (b) is the ground truth, image *X*′ (d) is the result of our method. Image *T*′ (e) is a direct combination of *A*′ and *B*′ using maximum composition. Fig. (f) shows more than 1% difference between *X* and *X*′ in red. In SSIM (g), the brighter are the images, the better. In optical flow (h), the darker the images are, the better. All scale bars are 200 μm.

### 3. Evaluation of final 3D models

To evaluate different approaches, we use nearest neighbor interpolation (a straightforward approach), truncate series to contain only “good” sections, utilize an optical-flow-based interpolation with alpha-blending (our method with blending per [67]), and use the method presented here, optical-flow-based interpolation with intensity-maximum-blending. All these methods are applied to the same spleen image series.

In all cases, the full 3D reconstruction pipeline is utilized. We start with a registered data set [68], apply normalization [89], color deconvolution [78], and above healing methods. After healing, inter-slice interpolation [67] is applied, followed by volume filtering. We apply 3D grayscale closing filter with radius 7 and a Gaussian blur with *σ* = 1. Then, the mesh is constructed with isovalue 120. We apply mesh repair [49] (octree depth 9), Taubin smoothing [103], and small component removal (at 2% of the main diagonal). Figure 1 visually compares the cropped and healed meshes. We then compute geodesic distances from the same starting point and over the distance of 100% of the main diagonal.

Figure 9 compares the features of the three complete meshes. The data sets are very dense and self-occluding. The difference between the methods is visible in the amount of connections in the blood vessels. In other words, where more interruptions are (due to missing or non-present repair), there more unconnected components ensue. Such components are colored in dark red.

**Figure 9:**
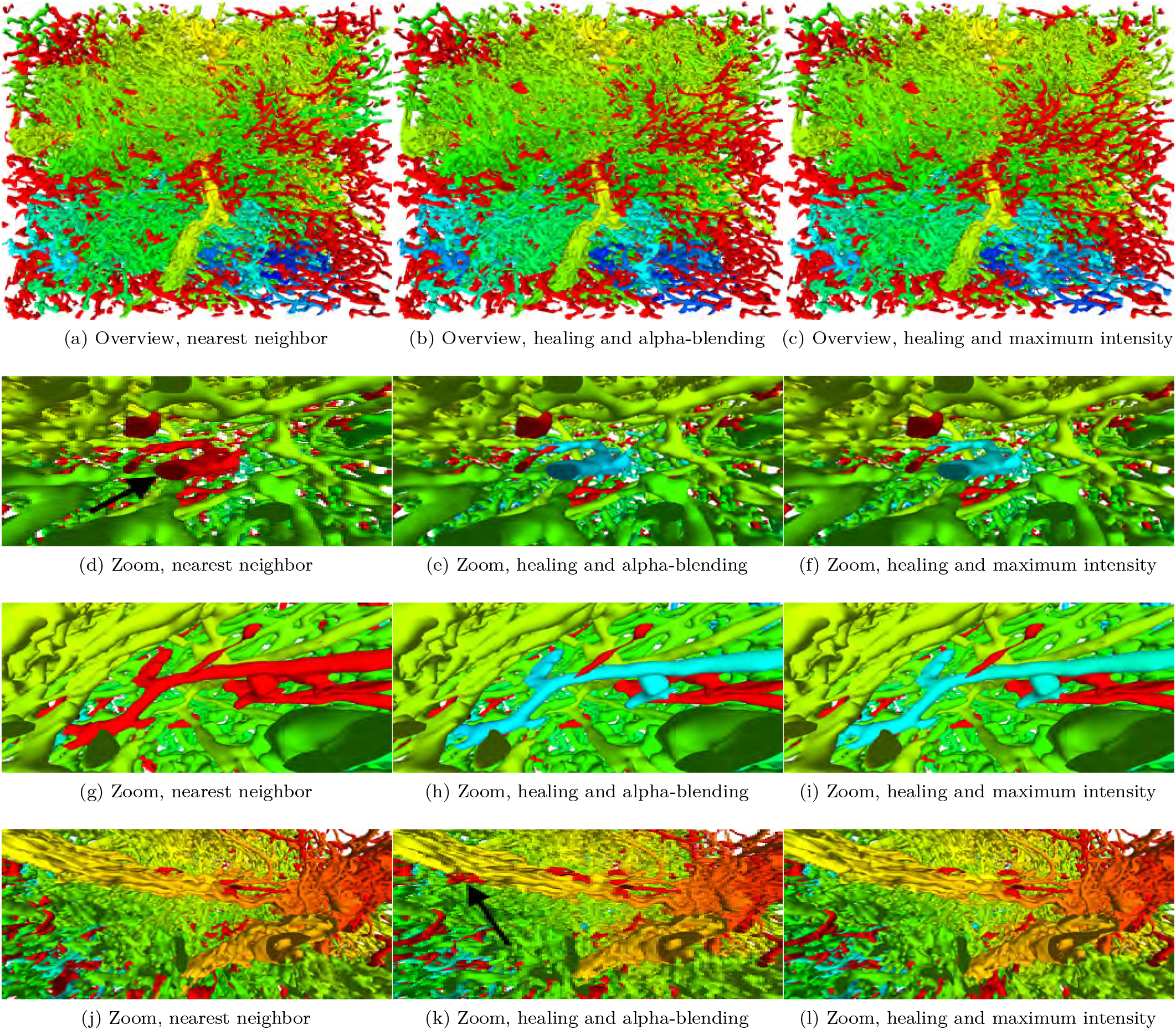
Geodesic distances from same origin for different kinds of healing applied to the same data set. Notice unconnected parts of blood vessels in dark red in other healing methods, marked with black arrows. Figs. (a)–(c) provide an overview, (d)–(l) show detailed views. The widths of the complete reconstructions (a)–(c) are 1 mm each.

If we zoom in (Figs. 9d–9l), we see some not-connected blood vessels in dark red in nearest neighbors and in alpha-blended meshes. Those blood vessels are connected with our method. To be more specific, some blood vessels were not connected with nearest neighbors (d), (g), or with optical-flow-based healing with alpha blending (k). Those blood vessels were connected when using optical-flow-based healing with maximum blending (f), (l). Supplementary video shows the mesh from (c) in motion.

We additionally evaluated all the vertices in those four meshes by their connectivity with above geodesic distance computation. Fig. 10 considers a) number of vertices having the geodesic distance of zero (i.e., not connected to the starting point) compared to the number of connected vertices; b) the violin plot [43] of geodesic distances for all vertices in each of the meshes. It appears hard to describe the differences between the reconstructions statistically.

**Figure 10:**
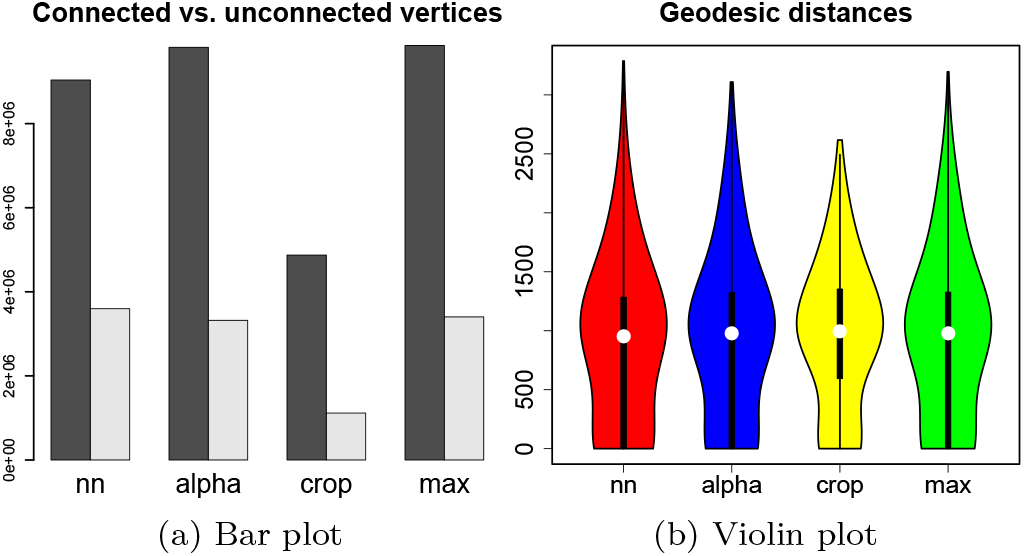
Statistical evaluation of reconstructed meshes. In (a), the dark bars show the number of connected vertices (the higher, the better), the light bars show the number of unconnected vertices (the lower, the better). The violin plots in (b) show the general distribution of the geodesic distances on all vertices. In both figures, “nn” stands for nearest neighbor, “alpha” for our healing method with alpha blending, “crop” for the truncated volume with no damaged sections, “max” for our healing method with maximum blending.

## IV. DISCUSSION AND FUTURE WORK

### 1. Existing methods

#### a. Registration

Registration methods typically assume a certain degree of similarity in the representation between the to-be-registered images. Focusing on registration of serial sections, in most cases the staining the same throughout the series (“unimodal” registration). In a classical multimodal registration the subject is the same, but the representations (i.e., the stainings) are different. A “common ground” needs either to be established with a segmentation, or similarities are established with a machine learning method. Basically, an intermediate representation, similar for all modalities is established. The registration operates on this intermediate, the actual transformations are then transferred to the original images.

In our case, the registration of an alternating series is a typical research task. Our stainings are established to answer a research question. This is a very different setting from establishing a multimodal registration for routine clinical stainings. Clinical stainings will be created in larger amounts for years after the establishing the registration. Hence, a machine learning approach makes economically sense, even with some additional manual labeling to bootstrap the learning. In our case, there are too many sections to be labeled by hand and too few to establish a proper machine learning method. Per design of the biological study, we have, however, a “common ground” in form of SMA and CD34 stainings in our data.

We previously used CD34 to establish a registration of uniformly stained serial sections. In the case of the registration of alternating series, the pure CD34 channel extraction by the means of color deconvolution did not work, but a denoised CD34 channel performed well.

Thus, we present here not an unconditional multimodal registration method that can operate on sections with no common ground via some similarity measure, but a fully automatic method to lift a registration method from uniform stainings to alternating stainings with a common ground.

Our contribution is thus different from all known to us related work on “unimodal” and multimodal registration. It is a very technical, but decisive and new contribution that allows for all the further processing in the 3D reconstruction pipeline. Our experience might be useful for others when designing a study with more channels than chromogens usable at once. We basically show the way for a conventional, “unimodal” registration (in our case: a fully automatic feature-based method) to “survive” an alternating stainings. The prerequisite is that some ubiquitous structures are present in both stainings, labeled in the same manner. Nerves or blood vessels are examples of such structures.

#### b. “Healing”

In the biomedical community a damaged histological section is often perceived as an inevitable artifact of lab work. Such artifacts are seldom reported in publications; basically a further, better section is to be produced to make the result publishable. This behavior is, however, not quite productive in 3D reconstructions from serial sections. A series needs to be consecutive for a 3D reconstruction. Should a section in the middle of a series be missing, only truncating a series or repair approaches are viable. A typical approach of others is to “repeat” the nearest neighbor section. Fig. 9 (d), (g) shows the drawbacks of this approach in form of broken connectivity in dark red.

All prior repair methods known to us cannot deal with damaged sections without loosing the connectivity of microvessels. Our method is able to maintain such a connectivity, which is novel.

In ref. [97] missing sections are reported to degrade the quality of registration. Our registration is able to bridge a missing section by repeating a neighbor. However, we had to introduce the presented method to maintain blood vessel connectivity. Burton et al. [21] have averaged nearest neighbor sections. We show that our method is more precise than the nearest neighbor repetition or even than the “healing” with alpha-blending (Fig. 9). Our method involves further non-linear distortions stemming from additional registration beforehand and from transformations induced by optical flow. Such non-linearity contrasts our approach with averaging. Our method is also more reproducible than sketching the missing sections from low-resolution unstained overview [91].

It appears to be viable to “heal” a missing section with existing mechanisms for the interpolation between sections [22, 67]. Indeed, we also used optical flow for such interpolations. However, we found that a correct blending method is crucial for the success of section repair, as Fig. 9k shows.

Summarizing, our method is more sophisticated and more precise in maintaining connectivity than other approaches. Our method is also more reproducible than the suggestions from the literature.

### 2. Evaluation criteria

It appears hard to us to precisely and practically evaluate the connectivity of the data in a very large, highly self-occluding data set. Using synthetic data can be an option, but real data sets are much more complex and might leave some cases unhandled in mock data. Image-based metrics (Tables I, II) show rather the correspondence of two images, whereas a good healing “imagines” a middle ground between two images. We found empirically that distorting the two neighbor images and then combining them to an intermediate produces better results, but this approach naturally slightly decreases the similarity between the intermediate and original neighbor images. It also appears that the repeated registration to *A*′ and *B*′ decreases the similarity between both distorted images as well. This is not bad as such, as long as the optical flow has enough common ground to perform well. Statistic evaluations (Fig. 10) of whole meshes are quite subdued: some broken-off parts might be a very minor issue statistically, but it would be appallingly clear during a visualization or a visual analytics session.

For the above reasons we regard a mesh-based evaluation of the repair methods crucial. In Fig. 9 we found some branches not connected in alternative methods that are connected in our suggested approach. A proper quality metric is also relevant in the context of the next section.

### 3. Deep learning

Image-based deep learning methods for recovery of missing image parts, e. g., ref. [115], are relevant and can be used to extend this work. Even more interesting are mesh- or voxel-based deep learning methods [16, 40, 47, 62, 88, 95, 114] with which a similar task to our original motivation could also be solved: interruptions in a 3D scene due to missing data could be detected, classified, and repaired. We further discuss this idea in the next section.

However, the big questions on the reproducibility of the results and on understanding of the actions of the network might hinder wider adoption of a deep-learningbased method in the area of medical imaging. Our healing method has a nice property of being a combination well-understood image transformations. Should a mistake in our method be detected, the source of the problem can be found easily.

Quality control criteria for connectivity repairs are quite easily defined. Corner cases are: something is not connected, but it should be; something is connected, but should not be.

### 4. Future work

Further adjustments of the registration technique are viable. These include experiments with further feature detection, description, and matching algorithms. An extension of matching to incorporate the structure of the data set could further improve the registration. Offloading more work to the GPU would improve execution times.

In general, registration methods for larger alternately-stained sections would be always sought for. The reason for this is that immunohistochemistry for transmitted light microscopy (and also, but less so, for fluorescence) is limited in the number of simultaneous “channels”, the number of antibodies that can be combined in a single staining.

Of course, we look forward to apply our healing technique to further series and organs. Optical microscopy images from serial semi-thin sections of animal lung probes look promising; Section III 2 b showcases our first attempts. A more confident investigation of human bone marrow stainings for blood vessels and certain progenitor cells becomes more viable. A healing technique becomes even more crucial in the context of larger human tonsil data sets, as this organ is often damaged during acquisition.

It still appears interesting to straightforwardly apply deep learning to image-based healing and to compare the result with our method. This study does not use machine learning methods by its design, as it aims for explainable intermediates. Future voxel- or mesh-based healing methods (potentially based on works mentioned above) might be instrumental in correction of the interruptions in biological mesh data.

## V. CONCLUSIONS

The contributions of this paper are twofold. Firstly, we found a way of registering alternating histological serial sections using an originally non-multimodal registration method. Secondly, we present a methodology to repair damaged or missing sections from a histological series.

With a simple, but previously not published trick, we registered a series of alternating serial sections, 148 images with dozens of thousands pixels per side. Our approach is based on color deconvolution and image filtering, followed by an existing feature-based registration. Our method is fully automatic. Basically, we have “lifted” a registration method that operated on a series with same stainings to a method operating on alternating series.

Damage happens routinely at all phases of processing serial sections, during biological processing, during acquisition, or even during digital processing—registration errors, for example. We were able to “heal” missing data from a series of sections. We reached our goal of recovering the lost 3D connectivity in a histological series with image-based methods. Our healing methodology promises the availability of larger series and larger potential 3D reconstructions, which facilitates better understanding of biological structures.

Specifically, applying the presented methods to a series of 148 immunostained serial sections, we were able to reconstruct more than 1 mm^3^ of a tissue at a microscopic resolution. In the same region only 84 sections could have been reconstructed with conventional methods.

## VI. ACKNOWLEDGMENT

The author is thankful to B. S. Steiniger (Philipps-University of Marburg) for the awesome possibility to work with digitized biological data that led him closer to the life sciences. Most of the biological data shown in this work originates from her lab. M. Guthe (University of Bayreuth) provided hardware, software, and his support. C. Mühlfeld (Hannover Medical School) supported the author with a position at the Hannover Medical School. A. Seiler, K. Lampp, and H. Pfeffer (Philipps-University of Marburg) were involved in the preparation and processing of the spleen serial sections. We thank A. Schauss (University of Cologne) for the acquisition of the spleen data. The author thanks R. Grothausmann, L. Knudsen, C. Mühlfeld (Hannover Medical School) and M. Ochs (currently Charité Berlin) for the availability of the lung data. S. Faßbender (Hannover Medical School) prepared the lung serial sections. M. Berthold (currently BCM Solutions GmbH) was involved in software development of the renderer used in the video. We used SSIM implementation by R. Mehdi.

This work was supported in part by German Research Foundation (DFG) grant no. 420759458. A part of this work was performed while the author was still at the University of Bayreuth.

## Notes

https:/doi.org/10.5281/zenodo.3544828

